# Optically Mapped Black Genomes: Distinct Structures and 22q11.2 Deletion Syndrome Mechanisms

**DOI:** 10.1101/2024.07.08.602568

**Authors:** Steven Pastor, Oanh Tran, Ryan Lapointe, Arnold Z. Olali, Douglas C. Wallace, Bernice Morrow, Elaine H. Zackai, Donna M. McDonald-McGinn, Beverly S. Emanuel

## Abstract

The genomic architecture of 22q11.2 Deletion Syndrome (22q11.2DS) has focused on analysis of white genomes. However, Black individuals appear to have a lower prevalence of 22q11.2DS compared to whites. To improve the understanding of different populations in relation to 22q11.2DS, optical mapping data from 106 genomes across various Black and white genomes were used to determine the organization of 22q11.2 genomic structures. This revealed extensive variability between the groups regarding copy number and orientation changes of the elements comprising the 22q11.2 low copy repeats (LCR22s). Several novel CNVs and whole haplotype configurations, private and of different prevalence to each group were detected. The diversity of CNVs within Black genomes compared to white genomes was especially striking. To determine the impact of this variability, Black families with *de novo* 22q11.2DS probands were compared to white families. The highly variable configurations of Black and white haplotypes led to several unique non-allelic homologous recombination (NAHR) scenarios with recombinations at different loci. In particular, Black families had unique recombinations yet to be observed. Thus, the unique and highly variable haplotype configurations of LCR22s in Black individuals may play a role in their decreased incidence of 22q11.2DS.

## 1. Introduction

The 22q11.2 deletion is the most common microdeletion syndrome with a prevalence of 1 in every 3-4000 live births [1,2,3]. Amongst those diagnosed with 22q11.2DS, it has been observed that Black patients are under-represented as compared to other ethnicities. To date, no genetic explanation for this has been described [4].

However, the lack of genetic exploration into the deletion between various patient groups is driven by the complexity of repeats within and flanking the region of the typical deletion. The 22q11.2 region contains eight Low Copy Repeats (LCR22s), labeled as LCR22A to LCR22H from centromere to telomere [5]. The LCR22s are comprised of sequence modules of varying lengths containing interspersed genes and pseudogenes [5]. Sequence analysis of these modules reveals a complex organization of duplicated elements [5]. In over 90% of 22q11.2DS cases, an approximately 3 Mbp de novo, hemizygous deletion is immediately flanked by LCR22A and LCR22D, two of the largest LCR22s [2,5,6]. Both LCR22A and LCR22D contain a highly homologous (>99% sequence identity) ∼160 kbp repeat sequence, termed SD22-4. This module is frequently present in more than one copy in LCR22A and occasionally in LCR22D [6,7,8]. The combination of large, near-identical segments makes the LCR22s substrates for non-allelic homologous recombination (NAHR), leading to genomic rearrangements. Unfortunately, these characteristics make the LCR22s difficult to sequence and to reliably identify rearrangement breakpoints in individuals with 22q11.2DS.

Optical mapping-based technologies have recently been employed to resolve the difficulty in producing whole haplotype maps and sequences of LCR22A and LCR22D [9,10]. Notably, LCR22A was found to be hypervariable between individuals, with variations in the number of copies and orientation of elements that comprise the LCR22s [9]. However, a comprehensive characterization of these structures within and between non-white populations of individuals and 22q11.2DS patients has yet to be performed. Since analysis of the 22q11.2 region has been dominated by genomes of European origin, there may exist bias in information about the structure of the region and ultimately genetic mechanisms which may play a part in NAHR. There is currently a need for population-level characterization of diverse genome sets to comprehensively understand the mechanisms across non-white populations in 22q11.2DS. [13].

Recent studies at the Children’s Hospital of Philadelphia (CHOP) determined that there is a paucity of 22q11.2DS diagnoses in Black patients that is not due to specific phenotypic differences. When controlling for congenital heart disease, the timing of 22q11.2DS diagnosis for Black and white patients at CHOP was similar. Further, co-morbidities were minimally different between the groups and craniofacial features were not a limiting diagnostic factor. Thus, it is hypothesized that 22q11.2DS is truly less common in Black patients and this may be explained by structural differences in the 22q11.2 LCR22s.

To resolve this conundrum, two comparative genomics operations were performed. First, the variability of the mapped structures in the 22q11.2 region were determined across 106 genomes encompassing several different populations of Black and white individuals. Second, previously mapped white and newly-mapped Black 22q11.2DS families were compared to one another and to the phenotypically normal individuals to find structures more frequently associated with NAHR. The goal was to determine if NAHR permissive structures are less prevalent in Black genomes, possibly explaining decreased occurrence of 22q11.2DS in Blacks. Overall, the study illustrates the advantage of using genomes from varying backgrounds to understand complex mechanisms in disease and represents a paradigm for future works in 22q11.2DS mapping and sequencing studies across populations.

## 2. Material and Methods

### Sample composition, collection, and consent

Subjects for the 22q11.2DS study were primarily families consisting of two healthy parents and an affected proband with a *de novo* ∼3 Mbp deletion in 22q11.2. The white cohort was comprised of 26 trios, one quad (identical twin probands), and three duos where the parent available was the parent of deletion origin (PoDO) (88 genomes). The Black cohort was comprised of three trios and two duos where the parent available was the PoDO (13 genomes). Patients and their non-deleted parents were tested for the presence or absence of the deletion using either a FISH assay with N25 probes (Abbot Molecular, Abbot Park, Illinois, USA) or the MLPA SALSA P250 DiGeorge diagnostic probe kit (MRC-Holland). Additional unaffected samples were collected from 40 unrelated whites who did not possess a deletion in 22q11.2. Informed consent was obtained from all participants and/or their legal guardians. They provided written consent for their EBV cell lines and DNA to be used for research purposes. The study was approved by the Children’s Hospital of Philadelphia under the Institutional Review Board (IRB) protocol 07-005352. All experiments were performed in accordance with the relevant IRB guidelines and regulations.

### Sub-Saharan African samples

Twelve unrelated, de-identified Sub-Saharan African lymphoblast cell lines were obtained with informed consent. These participants did not possess a deletion in 22q11.2. The transfer of these samples to Children’s Hospital of Philadelphia and continued study was approved under IRB protocol 10-007835.

### Coriell samples

Forty unrelated African Ancestry in SW USA (ASW) lymphoblast cell lines were procured from the Coriell Institute for Medical Research.

### Parent-of-deletion-origin detection

Fourteen short tandem repeat polymorphisms (STRPs) or microsatellite markers within the deleted region (see Supplementary Table S1 online) were used to determine the PoDO. Fluorescent modified PCR products were analyzed on an ABI 3730 instrument at the University of Pennsylvania Genomics and Sequencing Core. The GeneMarker software (Version 3.0) was used to analyze PCR fragments of an affected child and unaffected parents. An informative marker would typically consist of an affected child matching the fragment of one parent, but not the other. At least three informative markers per family were required to assign parent of origin.

### Ultra-high molecular weight (UHMW) DNA isolation for optical mapping

UHMW DNA was obtained following manufacturer’s protocol for gel plug lysis (Bionano, #30026, Rev. F), DNA isolation from fresh cells (Bionano, #30396, Rev. B), and fresh cell pellet DNA isolation (Bionano, #CG-0003, Rev. B). UHMW DNA was initially isolated by lysing fresh cells embedded in agarose gel but was later isolated by binding UHMW DNA extracted from fresh cells to a thermoplastic paramagnetic disk. Purified UHMW DNAs with high viscosity and a concentration of 36-150ng/μL were then labeled.

### DLE-1 labeling and chip loading

Purified UHMW DNA was labeled, purified again, and stained following two versions of Bionano’s Prep Direct Label and Stain (DLS) Protocol (Bionano, #30206, Rev. G and #30553-1, Rev. D). Labeled and stained samples with a concentration between 4 and 16ng/μL were loaded onto a dual flowcell Saphyr chip (Bionano) and imaged on a Bionano Saphyr instrument. After 24–36 h, a flowcell on a Saphyr chip can typically generate 1200-1500Gbp of data.

### Data pre-processing

Tab-separated BNX files containing molecule length, label quality scores, and label locations were output from the Bionano Saphyr optical mapping platform. BNX files were filtered for as long a length while maintaining the manufacturer’s recommended 320Gbp of total molecule data. Long molecules were required to uniquely span tandem duplicon modules, some ranging > 200 kbp, which is longer than the manufacturer’s default 150 kbp cutoff for molecules. Molecule qualities were assessed by the Bionano Access Molecule Quality Reports and compared to manufacturer’s recommended values (https://bionanogenomics.com/wp-content/uploads/2018/04/30223-Saphyr-Molecule-Quality-Report-Guidelines.pdf). Additional data were collected in samples not meeting these values.

### Localized *de novo* assembly pipeline

Human genome reference build 38 was *in silico* nicked with the direct labeling enzyme (DLE-1). Molecules from BNX files were mapped with higher confidence than default to each chromosome, except chromosome 22, using Bionano Genomics’ RefAligner (derived from Bionano Solve versions 3.3 and 3.4) program (notable parameters: -M 3 3 -FP 0.918057 -FN 0.099062 -sf 0.233588 -sd 0.090609 -S 0 -minlen 200 -minsites 15 - T 1e-25 -res 3.5 -resSD 0.7 -Mfast 0 -biaswt 0 -A 5 -BestRef 0 -nosplit 2 -outlier 1e-7 - endoutlier 1e-7 -RAmem 3 30 -hashgen 5 4 2.4 1.4 0.05 5.0 1 1 1 -hash -hashdelta 14 10 24 -hashoffset 1 -hashrange 1 -hashGC 300 -hashT2 1 -hashkeys 1 -hashMultiMatch 30 10). Chromosome 22 was not included in these alignments because of the high variability in LCR22 regions, potentially leaving out molecules otherwise mapping to legitimate haplotypes. The default molecule mapping p-value from RefAligner was increased to 1e-25 to ensure only higher confidence non-chromosome 22 mappings occurred and to ensure polymorphic chromosome 22-specific molecules did not map to another chromosome. The original BNX file was filtered for molecules mapping to all non-chromosome 22 maps using the RefAligner “skipidf” command.

Filtered BNX files were *de novo* assembled using default parameters in Bionano Solve v3.3 and v3.4. Molecules and Assembled consensus maps (CMAPs) were aligned to the *in silico* nicked hg38 reference map.

### Defining anchor and ambiguous regions in the hg38 reference map

Data of manually validated 22q11.2 haplotypes from 88 previously mapped genomes were used to define the sets of unambiguous DLE-1 labels required to span repetitive elements [9]. The hg38 reference coordinates of LCR22A and LCR22D were demarcated into frequently rearranging repetitive modules and less variable (based on DLE-1 label distribution) genomic regions. The regions exhibiting little to no variation in the 88 genomes were chosen as anchors flanking the segmental duplications in LCR22A and LCR22D and lacked segmental duplications themselves. Each anchor label set (>=12 DLE-1 labels and >=100 kbp) was defined as 18–18.15 Mbp for 5′ LCR22A, 19.035– 19.15 Mbp for 3′ LCR22A, 21–21.11 Mbp for 5′ LCR22D, and 21.565–21.7 Mbp for 3′ LCR22D. Molecules mapped to these regions do not map to other genomic loci without drastically changing the False Positive (-FP), False Negative (-FN), minimum mapped sites (-A), and minimum mapped length (-minlen) from the RefAligner parameters defined above.

### Paralogous label polymorphisms between SD22s to span ambiguous labels

The validated haplotype maps from the previously mapped 88 genomes indicated paralogous labels occasionally separating individual copies of SD22 modules [9]. Validated, assembled CMAPs were defined as containing molecules supporting connections between 5’ and 3’ anchored DLE1 labels and paralogous labels. In haplotype CMAPs lacking contiguous supporting molecules, manual curation was performed. Molecules were anchored and extended by following polymorphic labels between successive SD modules until the opposing anchor region was reached. A validated haplotype was defined as having >=5 molecules with at least 2 unambiguous labels (i.e., labels which do not appear in the opposing haplotype’s SD modules). All haplotypes and their unambiguously mapped molecules were manually confirmed using the Bionano Access visualization software v 1.4.1 (June 21, 2019 build).

### Confirmation of NAHR mechanisms

NAHR between LCR22A and LCR22D in PoDOs was determined through visualizing inherited LCR22 haplotypes in probands in Bionano Access. Label polymorphisms and label distance discrepancies present in PoDO haplotype maps compared to proband haplotype maps enabled the reduction of ambiguous recombination sites, as previously noted [9].

### Length Comparisons of Mapped Haplotypes

Starting and ending points in the LCRA maps were defined by the GRCh38 reference map 5’-most and 3’-most, respectively, DLE-1 labels residing only within segmental duplications. SD22-3 modules were defined as 200 kbp, SD22-4 as 160 kbp, and SD22-5 as 120 kbp. Five prime-truncated SD22-3 modules were defined as 100 kbp while three prime-truncated SD22-3 modules were 60 kbp. Consecutive modules were summed to generate lengths. The median of the minimum and maximum ranges for PoDO were used to disambiguate two haplotypes which hypothetically could undergo NAHR within the same genome.

### Statistical testing and Visualizations

Fisher’s exact tests, Chi-squared tests, and t-tests were performed in R version 4.4.0. Barcharts and boxplots were generated in ggplot2 v3.5.1. Haplotype map representations were created in matplotlib v3.9.0 with python v3.10.

## 3. Results

Optical mapping defines the hyper-variable landscape of SD22s across 106 genomes Optical maps of 106 genomes (212 complete haplotypes) across 7 groups of individuals were assembled and analyzed. There were 66 Black genomes and 40 white genomes. Haplotype maps of LCR22A through LCR22D were compared between the Black and white genomes to determine structural similarities or differences. Thus, the first step was to detect and analyze the baseline variation of SD22s in a population of phenotypically normal individuals, starting with LCR22A (Figure 1). The *in silico* labeled sequence of the GRCh38 reference is used as a baseline for orientation of genetic structures and to detect copies of 3 principal SDs, previously found to exist in LCR22A: SD22-3, SD22-4, and SD22-5 (Figure 1A) [9, 10]. Each of them was previously found to be copy number variable and orientation varied, between haplotypes (Figure 1B) [9, 10]. Further, positions of the modules may vary compared to the reference genome (Figure 1B, SD22-3 telomeric to SD22-4, which is opposite of the reference position). The distribution of all mapped samples is presented in Table 1.

**Figure 1.**
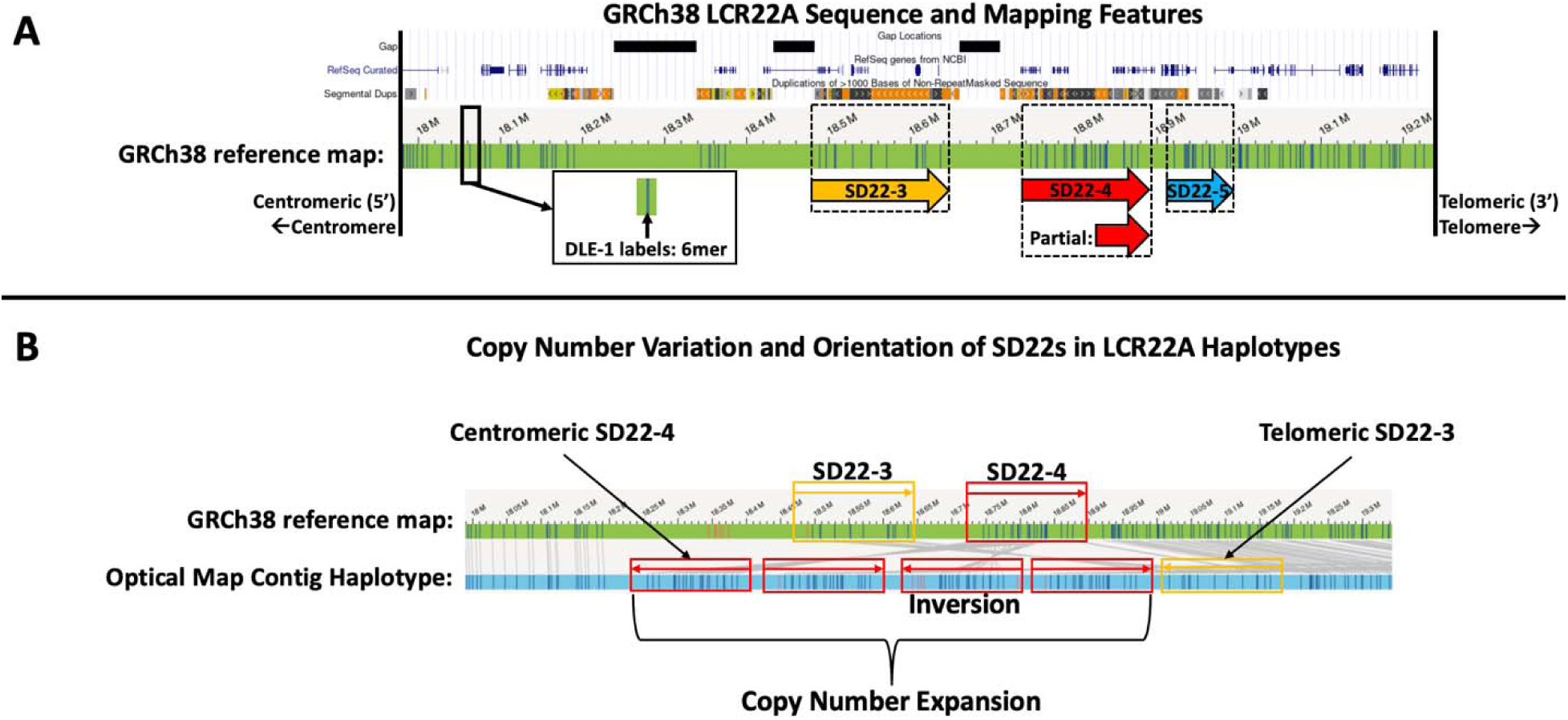
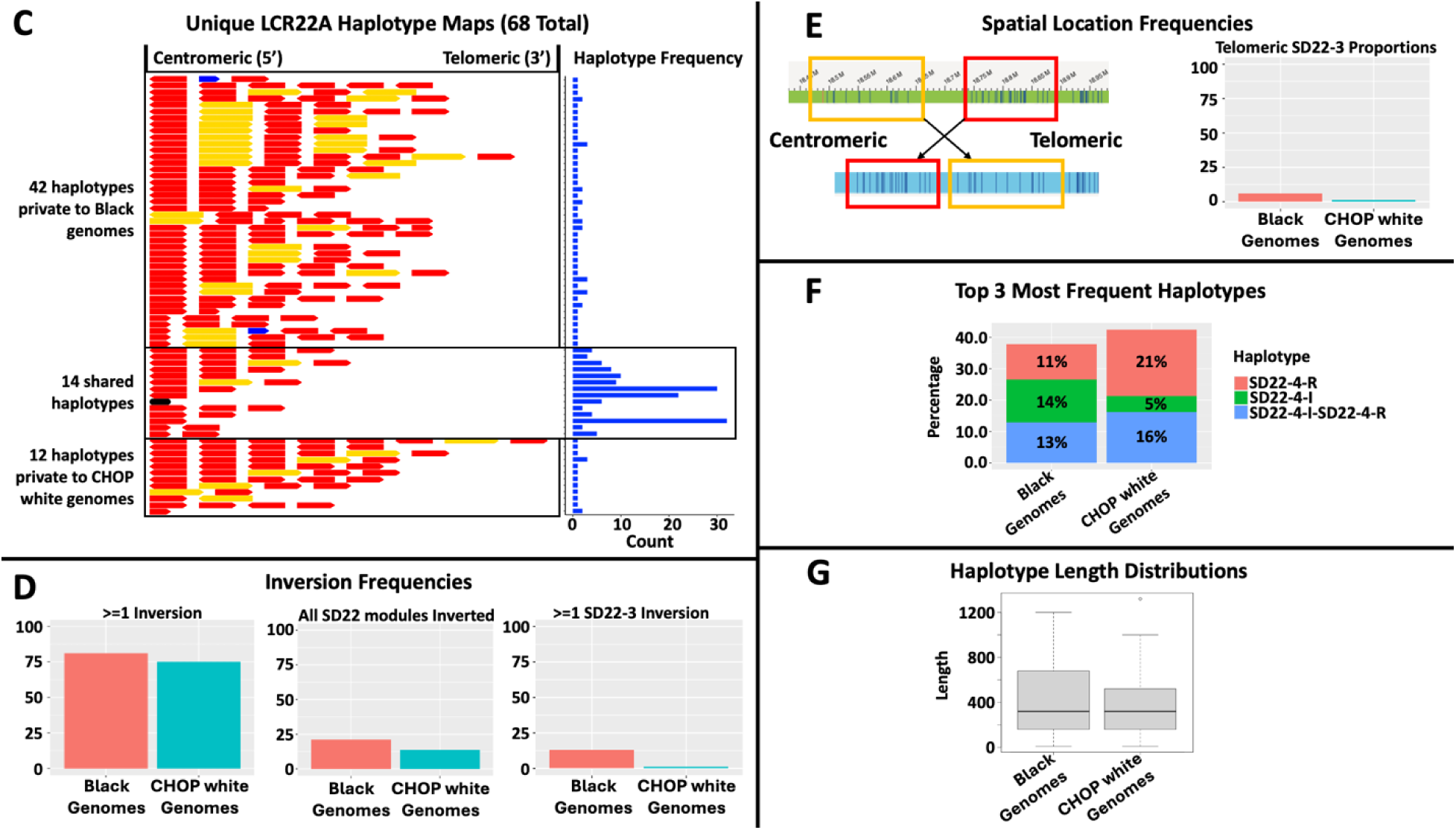
LCR22A haplotype map structures and length distributions. A. GRCh38 reference map (green) with DLE-1 labels (inset). Dashed boxes indicate 3 segmental duplication modules defined in previous studies (SD22-3, SD22-4 along with SD22-5). The 3 most frequent haplotypes trend toward fewer copy number increases and show that inverted SD22-4 is most frequent in Blacks while reference-orientation SD22-4 is most frequent in whites. its truncated representation, and SD22-5). Each of these modules are copy number variable (>=0 copies) and/or inverted relative to the reference. The SD22-4 (red) module was previously identified as extremely copy number and orientation variable between individuals, regardless of ethnicity or disease status [9, 10]. B. Example of copy number variation and orientation of LCR22A haplotypes, which dramatically differ from the GRCh38 reference and one another. Spatial location of SD22s is also variable, with any element able to “hop” around a haplotype (as shown with the haplotype’s SD22-3 as telomeric to SD22-4, which is opposite in the GRCh38 reference). C. Distributions of the LCR22A haplotype maps in phenotypically normal Blacks and whites. D. Black genomes have higher frequencies of inversions. E. Black genomes have higher frequencies of SD22-3 modules in the telomeric-most position. F. The 3 most frequent haplotypes trend toward fewer copy number increases and show that inverted SD22-4 is most frequent in Blacks while reference-orientation SD22-4 is most frequent in whites. G. No significant difference in lengths of haplotypes, indicating SD22-4 orientation and spatial location as the most important features between groups.

**Table 1.** Number of mapped, phenotypically normal, samples per group.

| Group | Number of Genomes |
| --- | --- |
| African Ancestry in Southwest USA (ASW) | 40 |
| African Caribbean in Barbados (ACB) | 4 |
| CHOP whites (CHW) | 40 |
| Esan in Nigeria (ESN) | 1 |
| Gambian in Western Division – Mandinka (GWD) | 6 |
| Mende in Sierra Leone (MSL) | 3 |
| Sub-Saharan Africans (SSA) | 12 |
| **Total** | **106** |

### Novel copy number variation and orientation of LCR22A SD22s in Black genomes

The 212 haplotypes produced 68 unique LCR22A haplotype combinations of SD22-3, SD22-4, and SD22-5 (Figure 1C). Of these, 43/212 configurations (20.3%) were present only once, with the remaining 25/212 (11.7%) in >=2 copies. There were 3.5x more haplotypes private to Black genomes (42) than whites (12), leaving 14 configurations shared between both groups.

Striking, was the high frequency of haplotypes containing at least one inverted SD22 module (Figure 1D). There are slightly more LCR22A haplotypes containing at least one inverted SD22 module in Black genomes (107/132 or 81.1%) compared to whites (60/80 or 75.0%). A subset of these configurations consists of all SD22 modules inverted. Black genomes contain more haplotypes with all SD22 modules inverted in LCR22A (21.2% vs 13.8% in whites, Chi-squared test p-value 0.2394). There are 15 heterozygous Black genomes with this configuration (22.7%), and 5 homozygous (7.6%). For whites, 12 are heterozygous (30%) and only 1 is homozygous (2.5%). The most striking difference in inverted modules was haplotypes with inverted SD22-3 modules. Blacks contained 17 haplotypes with at least 1 inverted SD22-3 module while whites had only 1 such haplotype (12.9% vs 1.3% in whites, Chi-squared test p-value 0.00714).

Another difference lies in the spatial location of SD22 modules. There is an increased frequency of telomeric SD22-3 in Blacks (6.1%) compared to whites (1.3%) (Figure 1E, Chi-squared test p-value 0.1827). Each of these haplotypes contains an inverted SD22-3 at the telomeric position. In contrast, the reference orientation SD22-4 module in the telomeric-most position appears more frequently in whites (80.0%), compared to Blacks (55.3%) (Chi-squared test p-value 0.00047). Further, Blacks have a slightly higher proportion of haplotypes with reference orientation SD22-4 modules in the centromeric position (29.5%) while whites have more of the inverted SD22-4 at that position (26.3%).

Three other configurations present in LCR22A were analyzed (Supplemental Figure 1). First, the LCR22A haplotype devoid of SD22-3, SD22-4, or SD22-5 appears once in white genomes (1.3%) and 4 times in Blacks (3.8%). The SD22-5 module appears twice, each time in a different Black genome. Finally, truncated SD22-4 modules appear at 7.5% in whites and 10.6% in Black genomes.

The 3 most frequent LCR22A complete haplotypes across all genomes are those with one reference-orientation SD22-4 (“SD22-4-R”), one inverted SD22-4 (“SD22-4-I”), or those with a centromeric inverted SD22-4 followed by a reference-orientation telomeric SD22-4 (“SD22-4-I-SD22-4-R”). Of note, Black genomes had over twice as many “SD22-4-I” haplotypes (13.6%), compared to whites (5.0%) (Chi-squared test p-value 0.07733) (Figure 1F). White genomes contained a higher frequency of the “SD22-4-R” haplotype (21.3%) compared to Black genomes (11.4%) (Chi-squared test p-value 0.07992). Finally, the “SD22-4-I-SD22-4-R” haplotype was similar in both groups (16.3% in whites and 12.9% in Black genomes). These results indicate that blacks have a higher frequency of inverted SD22-4 modules, albeit at non-significant amounts.

Despite the spatial and structural differences of LCR22A haplotypes between groups, the difference in lengths of LCR22A haplotypes between Blacks (mean = 436,136bp) and whites (mean = 399,125 bp) is not significant (Figure 1G; mean: Welch t-statistic, p-value = 0.3229) and the medians are exactly the same at 320,000 bp. Interestingly, the shared haplotypes between the 2 groups (see Figure 1C) are significantly shorter (<=4 copies of SD22s and mean = 357,857 bp) than haplotypes unique to either group and contain less copy number variation (all unique vs shared: Welch t-statistic, p-value = 6.71e-06, White unique vs shared: Welch t-statistic, p-value = 0.004751, Black unique vs shared: Welch t-statistic, p-value = 6.948e-06). The LCR22A haplotypes private to Blacks (mean = 710,000 bp) were of similar length to those in whites (mean = 708,333 bp). Thus, the principal differences between groups in LCR22A are derived from SD22 spatial positioning and orientation of SD22-4 and SD22-5 modules, specific SD22 copy number, and the presence of 2 other SD22s; SD22-3 and SD22-5. These 2 modules appear only in haplotypes unique to either group.

### Black genomes exhibit higher frequencies of inversions in LCR22D

LCR22D has less copy number variation than LCR22A, confirming previous results for white as well as Black genomes (Figure 2) [9, 10]. The major SD22 module present in LCR22D is SD22-4, which shares >99% sequence identity with the SD22-4 in LCR22A and is defined as the directly-paralogous-LCR22 (DP-LCR22) predominantly involved in NAHR [9, 10]. The GRCh38 reference map is used as a guide for copy number and orientation of the SD22-4 module and a frequently observed inverted sequence (Figure 2A). As in LCR22A, there is a greater frequency of full-length or truncated SD22-4 inversions in Blacks (15.9%) compared to whites. No white LCR22D haplotypes contain inverted SD22-4 modules (Figure 2B). The only major variation in whites are 3 genomes with a heterozygous haplotype containing a truncated reference orientation SD22-4 duplication, which is also contained in 3 Black genomes (top-left, Figure 2B). Thus, more SD22-4 inversions are present in Black genomes. Since SD22-4 is the predominant DP-LCR22 involved in NAHR, this potentially has implications for NAHR frequency in Black genomes.

**Figure 2.**
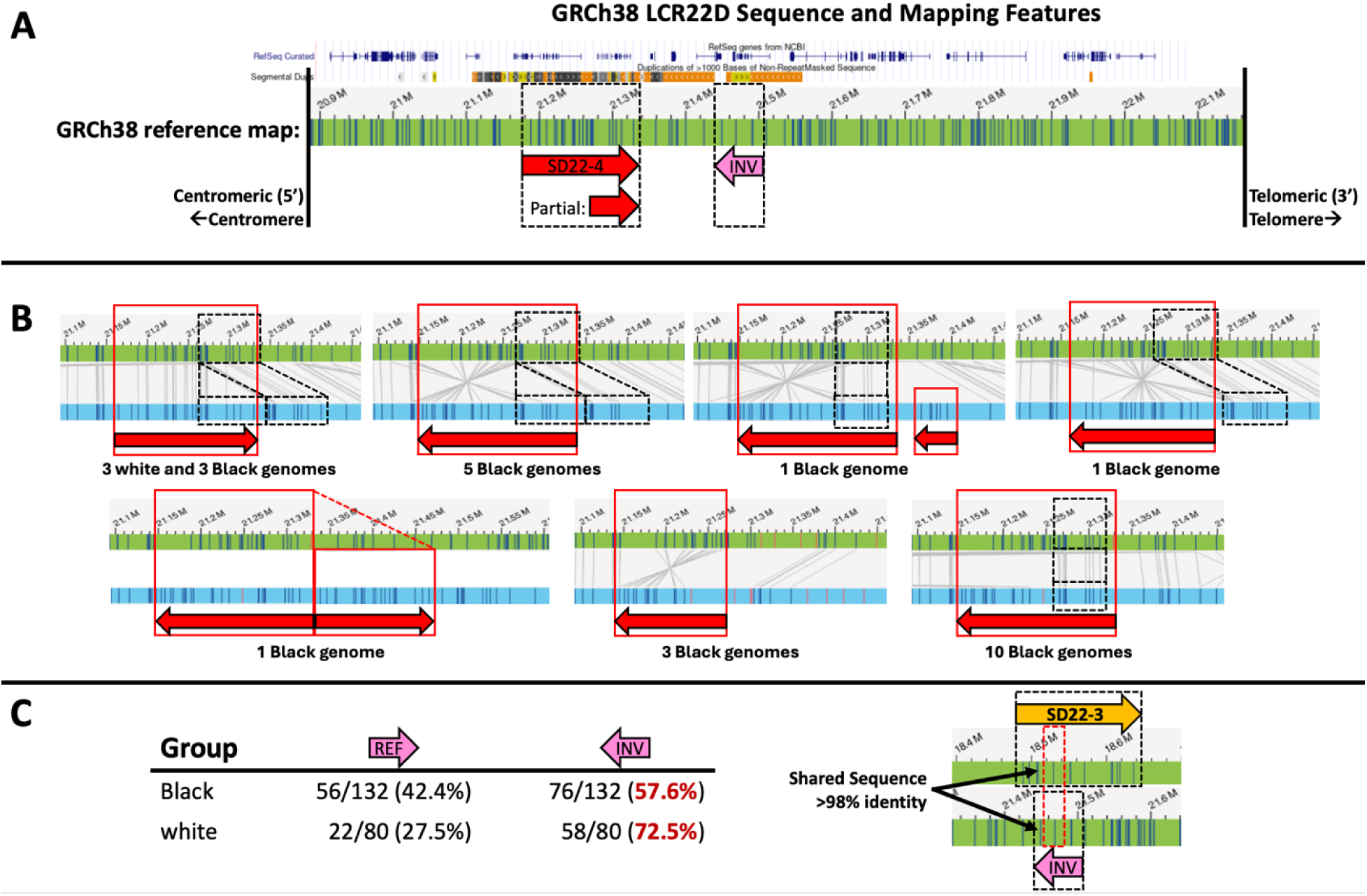
LCR22D haplotype map structures and length distributions. A. GRCh38 reference map for LCR22D. The 3 primary modules of interest are the fully-intact SD22-4 (red) and its partial variant along with the highly frequent, smaller inverted module (pink). B. Landscape of inversions and truncated duplications in LCR22D. Only Black genomes contained inverted fully-intact or truncated SD22-4 modules and these were typically accompanied with truncated duplications. One instance of a fully duplicated SD22-4 was found, and this haplotype contained an inverted SD22-4 followed by telomeric, reference orientation SD22-4 (bottom-left). Whites only contain one principal variant in the form of a truncated duplication of SD22-4 (top-left). C. The frequent ∼64 kbp inversion appears more frequently in whites than Blacks and shares sequence identity with SD22-3. C. Black genomes have lower frequencies of a ∼64 kbp inversion.

One notable inversion was previously described and is present in high frequency across all genomes, regardless of background (GRCh38 optical mapping coordinates chr22:21,448,904-21,491,649 (42,745 bp), Figure 2A, pink inverted arrow labeled “INV”) [10]. Approximately 72.5% of whites contain this inversion compared to approximately 57.6% of Blacks (Figure 2C, Welch t-statistic, p-value = 0.04162). This is the only inversion in either LCR22A or LCR22D seen more frequently and significantly in white genomes. A ∼33 kbp sequence of this module shares >98% sequence identity with 18,518,837-18,552,128 of SD22-3.

Finally, the LCR22B and LCR22C regions are much shorter in length than LCR22A and LCR22D, and exhibit far fewer variations between the 2 groups (Supplemental Figure 2). Since the results here are focused on the typical, ∼3 Mbp deletion between LCR22A and LCR22D and the DP-LCR22s directly involved in its inception, the remaining analyses only focus on LCR22A and LCR22D.

### Comparing the landscape of NAHR in Black and white parents-of-deletion-origin

There are 30 white PoDOs and 5 Black PoDOs available for comparison. As described previously, to determine a NAHR haplotype, paralogous labels can be used to delineate the boundaries of LCR22A and LCR22D in the deletion-containing haplotypes of 22q11.2DS probands [9, 10]. Figure 3A generally outlines this strategy, which is applied to the 35 deletion haplotypes of interest (see Supplementary figure 2 for all 35 scenarios). Since the LCR22A haplotypes are so variable between genomes, typically only the proband and the PoDO are required to determine NAHR haplotypes. Here, trios are used in 28/30 instances (88 genomes) for white families and 3/5 instances (13 genomes) for Black families.

**Figure 3.**
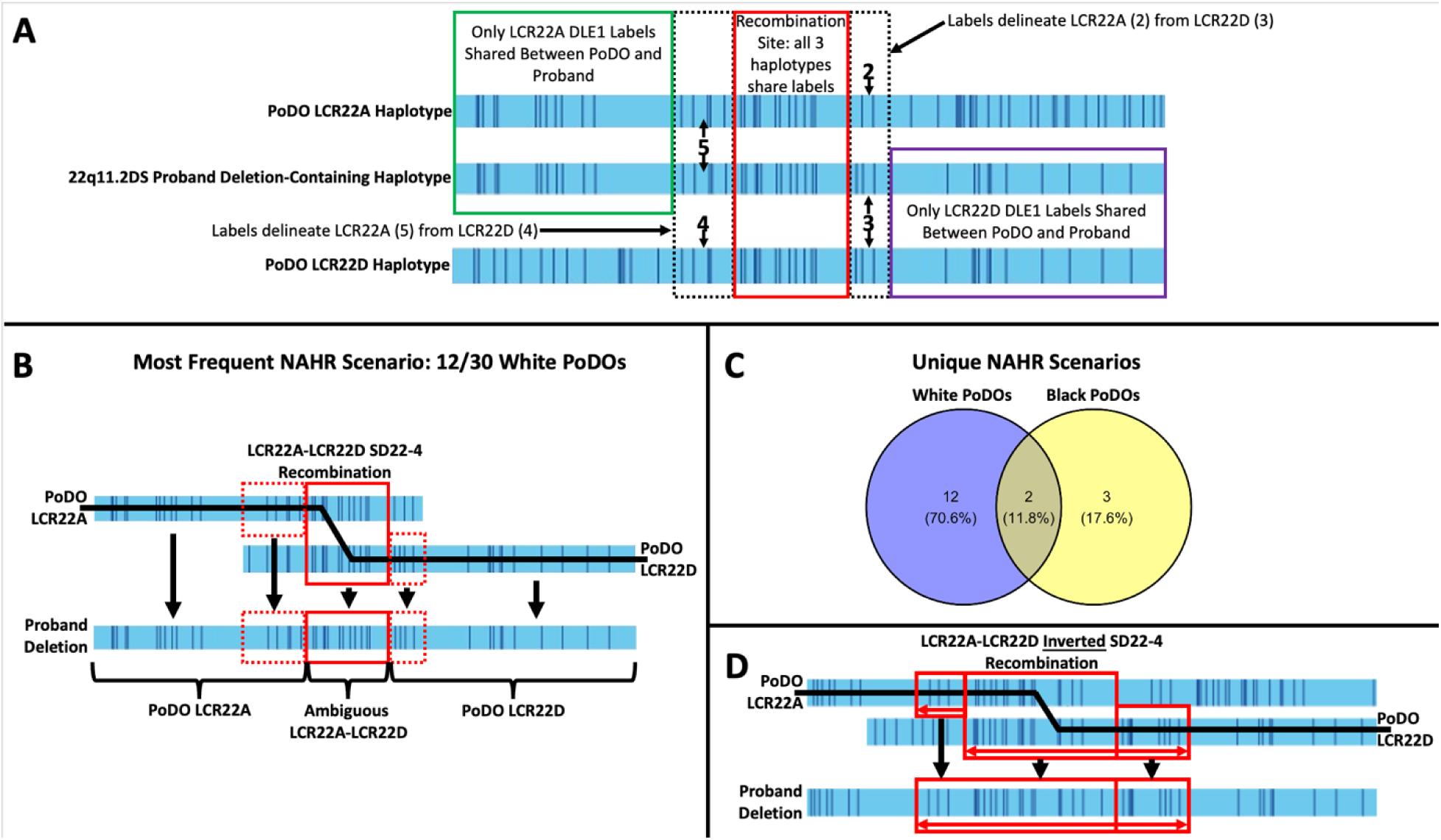
Landscape of NAHR scenarios in Black and white genomes. A. Strategy for delineating LCR22A and LCR22D modules in NAHR haplotypes from 22q11.2DS proband deletion-containing haplotypes. From top-to-bottom, comparing PoDO LCR22A labels to proband deletion-containing labels and to PoDO LCR22D labels enables delineation of NAHR haplotypes and sites of recombination by discerning shared and unshared labels. **B.** Most frequently occurring NAHR scenario in white PoDOs is only observed in white PoDOs. Normal white genomes contained this exact LCR22A haplotype in the highest frequency (21.0%) along with this LCR22D haplotype (72.5%), thus indicating a direct relationship of haplotype frequency in the general population and NAHR haplotypes, per population. **C.** Unique NAHR scenarios shared between the Black and white PoDOs indicates most scenarios private to either group and population-specific due to vastly different LCR22A and LCR22D haplotypes. D. The only NAHR scenario involving an inverted SD22-4 module in both LCR22A and LCR22D occurs in a Black parent.

To understand the structure of SD22s in the context of NAHR, the landscape of NAHR across Black and white PoDO genomes was analyzed. In white PoDOs, one scenario is observed as the most frequent (Figure 3B). Here, 13 of the 30 (43.3%) LCR22A NAHR haplotypes contain a single, reference orientation SD22-4 module. In the 13 NAHR events consisting of this haplotype, the LCR22A SD22-4 module mis-aligns with its DP-LCR22, the reference orientation LCR22D SD22-4 module. Interestingly, the most prevalent LCR22A haplotype in normal white genomes is one with a single, reference orientation SD22-4 module, which directly translates into the most frequent configuration leading to NAHR. Of these, 12/13 occur within the boundaries defined as SD22-4. The one genome where it occurs outside SD22-4 has the site of aberrant recombination in a paralogous sequence, centromeric to the SD22-4 module (GRCh38 coordinates: 21,101,987-21,137,158). This sequence module appears once in LCR22A SD22-4 and shares >99% identity.

Not all NAHR events require LCR22A and LCR22D reference orientation SD22-4 modules to align (Supplemental Figure 3). There are 7 events which do not involve direct alignment of LCR22A and LCR22D SD22-4 modules. Interestingly, one event involves an inverted LCR22A SD22-4 module misaligning with a portion of a reference orientation LCR22D SD22-4 module. Another event of note involves a portion of the reference orientation SD22-3 module (GRCh38 coordinates: 18,496,729-18,508,639), which also shares the aforementioned sequence with SD22-4 at >99% identity, indicating a repeated recombinogenic sequence (GRCh38 coordinates: 18,754,039-18,765,960). This was previously identified as long intergenic non-coding RNA (lincRNA) family *FAM230* sequence [9, 10]. For notable spatial organizations, 24/30 (80%) of events involve misalignment of a telomeric LCRA SD22-4 module (Supplemental Figure 4). Finally, comparing PoDOs to normal white and Black genomes, there are only 4 NAHR haplotypes which are private to PoDOs, indicating that whole haplotype configurations are unlikely to be the prime determining factor in driving NAHR. There is a linear relationship between frequency of a haplotype in the white population and its participation in NAHR of white PoDOs (Pearson’s Correlation Coefficient = 0.910914 at p-value = 4.606e-13 for the comparison between white normal haplotype frequencies and white PoDO haplotype frequencies. Black genomes did not exhibit this same correlation but the small sample size of PoDOs makes this comparison difficult (Pearson’s Correlation Coefficient = 0.1298028 at p-value = 0.3359 for the comparison between Black normal haplotype frequencies and Black PoDO haplotype frequencies).

Only 2 of the 5 Black LCR22A NAHR haplotypes are also present in whites but only 1 of the 2 has the same recombination locus (Figure 3C). Further, only 1 of the LCR22A NAHR haplotypes is private to Black PoDOs compared to white PoDOs or either normal genome group. This again signifies that whole haplotypes are not always indicative of NAHR events. For spatial positioning, 2 of the 5 (40%) scenarios involve telomeric reference orientation LCR22A SD22-4 modules. This is half the rate it occurs in white PoDOs, albeit in a much smaller sample size. For inverted modules, one scenario involves an inverted SD22-4 module and another involves an inverted SD22-3 module, which is not observed in white PoDOs. One of the most frequent haplotypes in normal genomes is the single copy reference orientation SD22-4 in LCR22A but with a nearly 2-fold higher proportion in white versus Black genomes. In contrast to whites where 12 NAHR scenarios involve this haplotype mis-aligning with a DP-LCR22 SD22-4 in LCR22D, no Black PoDO has this configuration. In fact, every Black NAHR LCR22A haplotype had >=3 copies of SD22s. Overall, with 2 never-before-observed NAHR LCR22A haplotypes and 4 completely different haplotype-recombination sites, Black PoDOs, even at a small sample size, have novel recombination mechanisms compared to whites.

In LCR22D, every white NAHR involves a fully-intact LCR22D SD22-4 module directly reflecting the LCR22D haplotypes observed in normal white genomes. Despite this, the actual sequences directly participating in NAHR may be within this module or in other paralogous loci. These are paralogous sequences which appear multiple times in LCR22A and LCR22D and have been previously described [9, 10]. The only NAHR event of note from whites is one with a haplotype containing a fully-intact SD22-4 module followed immediately by a truncated SD22-4 module (Supplemental Figure 5). This haplotype is also observed in 3 normal white and 3 normal Black genomes, indicated in Figure 2. The fully-intact SD22-4 module is where the site of recombination occurs, continuing the trend of full SD22-4 modules of LCR22D participating in LCR22D NAHR haplotypes of white genomes. The LCR22D recombination region ranged anywhere from 21,093,033-21,359,141, with 24/30 recombinations occurring due to DP-LCR22s via SD22-4 modules. All these whole LCR22D haplotypes appear in either normal white or Black genomes.

In Blacks, 4 of 5 NAHR events involve fully-intact reference orientation LCR22D SD22-4 modules. One contains an inverted truncated SD22-4 module and the recombination occurs directly within this inverted module (Figure 3D). This is not observed in any white PoDOs. In fact, no white PoDOs have an inverted SD22-4 module in LCR22D, regardless of the haplotype being involved in NAHR. Normal Black genomes are the only ones with inverted SD22-4 modules in LCR22D so this again reflects a group-specific NAHR event. Aside from this difference, the same recombination range and all haplotypes, are shared with white PoDOs.

### Factors driving NAHR differentials in Black and white genomes

It is hypothesized that there are likely two driving forces behind the lower prevalence of NAHR events leading to 22q11.2DS in Black families; the length of the nearly identical recombining sequences between LCR22A and LCR22D and distance between recombining elements from LCR22A to LCR22D [12]. Since LCR22 haplotypes private to either Black or white genomes are near identical in length, one hypothesis for 22q11.2 NAHR permissiveness may lie in the spatial orientation of individual SD22 modules rather than copy number.

### SD22 modules’ influence on contiguous sequence identity between DP-LCR22s

The first driving force seems to be that the longer the sequence identity between LCR22 modules the more likely the misaligned sequence will provide a site permissive for NAHR. From optical maps, sequence from GRCh38 and the CHM13 single haplotype sequence, the longest DP-LCR22 between LCR22A and LCR22D is the SD22-4 module. Thus, LCR22A haplotypes with all SD22-4 modules inverted relative to the LCR22D reference orientation SD22-4 module(s) or vice versa, would lead to shorter contiguous recombinogenic sequences between LCR22A and LCR22D (Figure 4A). It is observed that 22/30 NAHR scenarios in whites and 3/5 in Black PoDOs contain DP-LCR22 (SD22-4) modules directly involved in the recombination event leading to a transition from LCR22A to LCR22D. Previous work has defined this mis-alignment as approximately 160,000 bp [8]. Given the SD22-4 modules are the longest shared sequences between LCR22A and LCR22D, it is no surprise that the 8 events lacking DP-LCR22s contain shorter lengths of homology upon mis-alignment (20-60 kbp). Black PoDOs have the only event involving an inverted SD22-3, although with fewer samples. Both groups have one event with inverted SD22-4 in LCR22A participating in NAHR. The approximate average NAHR homology, based on map labels, is 127 kbp for whites and 130 kbp for Blacks, indicating a similar length of homology between the groups. It is important to note that these scenarios are only relevant if all modules are inverted in only one LCR22 and not both (e.g., only LCR22A or only LCR22D), otherwise inverted modules in both LCR22A and LCR22D become DP-LCR22s. Thus, both groups contain higher frequencies of events with long homology via DP-SD22-4 modules and with similar ranges of contiguous sequence identity.

**Figure 4.**
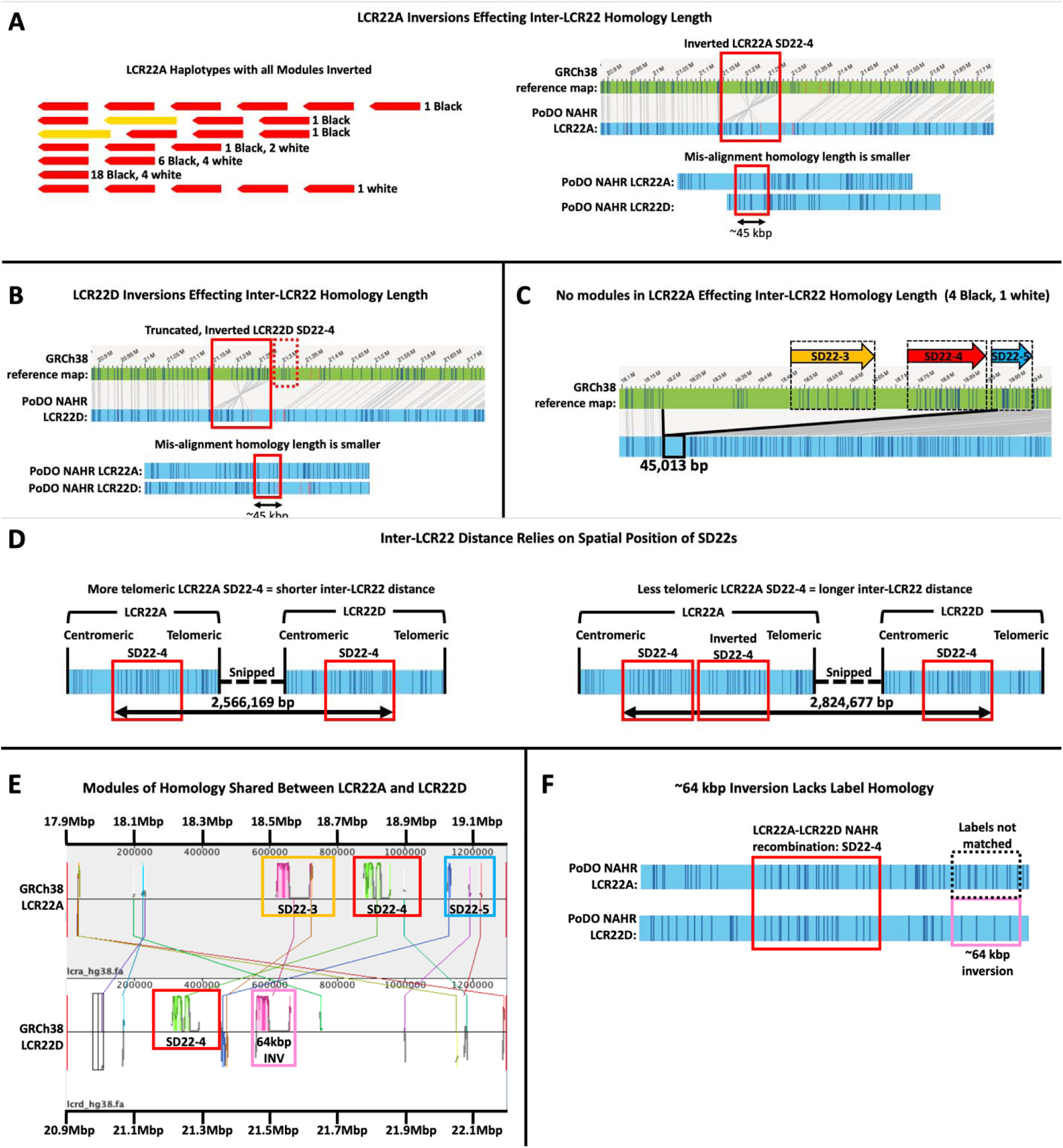
Driving forces analyzed in NAHR. A-C. Inversions and absence of SD22 modules lead to shorter mis-alignment homologies between LCR22A and LCR22D in NAHR. A. Normal Black genomes contain more LCR22A haplotypes with all SD22 modules inverted (left). This leads to shorter lengths of homology, assuming mis-alignment with reference-orientation SD22-4 in LCR22D (right). B. Only Black normal genomes contain LCR22D haplotypes with all SD22 modules inverted (top). This leads to smaller lengths of homology, assuming mis-alignment with reference-orientation SD22-4 in LCR22A (bottom). C. Four normal Black normal genomes contain no SD22 modules in LCR22A compared to only 1 white genome. This leads to a shorter length of homology for NAHR but no NAHR event with this haplotype is observed. D-G. Spatial location, especially telomeric modules, affects inter-LCR22 distance in NAHR. D. The inter-LCR distance between LCR22A and LCR22D SD22-4 modules was 2,566,169 bp if considering the centromeric DLE-1 label in LCR22A SD22-4 and the telomeric DLE-1 label in LCR22D SD22-4. Centromeric SD modules were rarely observed, as this distance would increase, leading to less mis-pairing. NAHR between telomeric reference-orientation SD22 modules has a shorter inter-LCR22 distance and may promote NAHR. Distance increases via telomeric inversions, which are more prevalent in Black genomes; however, Black PoDOs contain 3/5 events in non-telomeric SD22s, indicating population-specific haplotypes influence population-specific NAHR events. E. All possible contacts between LCR22A and LCR22D. F. The ∼64 kbp inversion may not promote NAHR at the DLE1 label level due to lack of homologous labels in recombinogenic segments of LCR22A and LCR22D.

Since DP-LCR22s offer the longest homology and there is a higher frequency of white and Black PoDOs with DP-LCR22s as the site of NAHR, LCR22A or LCR22D whole haplotypes with every SD22 module inverted decreases the length of homology. As mentioned, Black genomes contain a higher frequency of all-inverted LCR22A haplotypes, albeit with a large p-value (Chi-squared test p-value 0.2394). Perhaps the largest difference lies in LCR22D, where only Black genomes contain inverted SD22-4 modules, reducing the length of homology with LCR22A (Figure 4B). The LCR22D inversions participate in NAHR, but only one Black PoDO. Given the lower frequency of NAHR events involving inverted SD22-4 modules, it is possible such configurations reduce the propensity for NAHR to occur. Combining this with the higher frequency of all-inverted haplotypes in Black genomes could help explain the decreased incidence of NAHR in Blacks.

Also, supporting the first driving force is the absence of major SD22s in some LCR22A haplotypes, eliminating DP-LCR22s between LCR22A and LCR22D (Figure 4C). If the putative main recombination is between LCR22A and LCR22D SD22-4, a haplotype lacking this module would have opportunities to recombine but contain significantly smaller stretches of sequence identity. At a length of 45,013bp, this is much smaller thanthe ∼160,000bp afforded by mis-alignment of SD22-4 modules. Haplotypes of this sort are not found in any NAHR-associated haplotypes. Black genomes have the most with 4 such haplotypes and whites have only 1. White PoDOs had 1 such haplotype but this was a haplotype containing structures not participating in the recombinogenic process of NAHR. Again, Black genomes contain haplotypes with presumably less permissive NAHR structures than whites, but these haplotypes occur with less frequency than those containing any combination of >=1 SD22 module(s). Overall, Black genomes contain more inversions and other structures which reduce configurations favorable to NAHR, assuming length of homology is a driving force.

### Telomeric reference-orientation SD22 modules have decreased incidence in Black normal genomes and PoDOs revealing new mechanistic insights in Black PoDOs

The second driving force is based on the inverse relationship between NAHR and inter-LCR22 distance. The longer the distance between recombinogenic sequences the less likely the incidence of NAHR [12]. In support of this is the observation that the majority of observed NAHR mis-pairings involve the more telomeric SD22 modules in LCR22A (Figure 4D). The orientation of SD22s within the haplotypes, especially in the form of inversions, may lead to differences in NAHR frequency between Black and white genomes (Figure 4D, bottom scenario, indicating inverted SD22-4 blocking NAHR to the more centromeric SD22-4). In white genomes, 24/30 (80%) NAHR events involve an SD22 in the telomeric-most position of LCR22A, which directly participates in the mis-alignment. In normal whites, 80% of LCR22A haplotypes have SD22-4 telomeric modules in reference orientation (Figure 4B). Interestingly, Black PoDOs have only 2/5 (40%) NAHR events like this, and normal Black genomes have a decreased incidence of these haplotypes as well (55.3%). These results lead us to 2 conclusions; 1) the decreased incidence of telomeric reference-orientation SD22-4 modules in Black genomes leads to decreased frequency of NAHR in Black individuals or 2) the decreased presence of telomeric reference-orientation SD22-4 modules has less influence on NAHR in Black genomes. With more samples, it will be possible to test this further.

Due to the variable rearrangement scenarios derived from white and Black PoDOs, it remains clear of the possibility of different haplotype configurations and SD22 structures which are the direct sites of recombination. Thus, while inter-LCR22 distance and homology between DP-LCR22s, especially the SD22-4 modules, are likely driving factors in NAHR, the sheer number of other modular identities also clearly drives NAHR (Figure 4E). If these structures are present in the general population and are translated to the PoDOs, this leads to population-specific recombination configurations. Further, the possibility of inverted elements observed in Black PoDOs, and the higher presence of inverted elements in normal Black genomes may indicate other mechanistic structures which can recombine. Thus, by evaluating the Black PoDOs and normal genomes, it was found that inverted and non-SD22-4 elements may participate in NAHR more than imagined by solely evaluating white genomes alone. While the 2 driving forces’ influences on the decreased incidence of NAHR in Black genomes may be currently inconclusive, at the least, these results indicate that analyzing only white PoDOs may not be enough to draw conclusions about the likelihood of NAHR in Blacks.

### LCR22D inversions may promote decreased frequency of NAHR mis-pairings in Black genomes

The last major difference between groups is the presence of a frequently-inverted ∼64 kbp module in LCR22D, found to be significantly higher in white genomes (72.5%) than Black genomes (57.6%). White PoDOs also contain a high frequency of this module, at 78.3% and interestingly, so do Black PoDOs at 80.0%. In NAHR haplotypes, white PoDOs had this module in 26/30 haplotypes (86.7%) while Black PoDOs had 5/5 (100.0%). There is a significant difference between all 4 groups overall (Welch’s ANOVA one-way analysis of means, P-value = 0.01333) but pairwise comparisons revealed only Black normal genomes and white PoDOs as significantly different (0.026, Bonferroni-adjusted P-value of pairwise t-tests with pooled SD). When looking at both the GRCh38 optical map and reference sequence for the putative recombinogenic sequences between LCR22A and LCR22D upon mis-alignment, only instances of interspersed small sequences (<700 bp) covering 25% of the LCR22D 64 kbp inversion sequence may pair and each pairs with less than 85% identity (Figure 4F). Thus, it is unclear whether this module would explain fewer NAHR instances in Black genomes.

Finally, a rough approximation of the inter-LCR22 distance is calculated, using the centromeric DLE1 labels starting the recombinogenic modules in NAHR for each PoDO and ending in telomeric DLE1 labels (see distribution of inter-LCR22 distances, Supplemental Table 1). On average, Black PoDOs had a 2,680,355 bp average inter-LCR22 distance in NAHR events while white PoDOs were shorter at 2,612,545 bp. The decreased distance is due, in part, to the telomeric modules in LCR22A participating in NAHR in whites. Thus, inter-LCR22 distances in Black PoDO NAHR events, albeit with small sample size, are longer than in white PoDOs. Given the smaller number of telomeric reference orientation SD22-4 modules in Black genomes, it is possible that nested modules may participate more frequently in Black PoDOs and inter-LCR22 distance is increased. Again, this may help explain the decreased frequency of NAHR events in Blacks or it may be a feature in Black NAHR events.

## 4. Discussion

Chromosome 22, particularly region 22q11.2, is one of the remaining complex regions in the human genome left to be resolved due to its highly repetitive nature [11]. To date, optical mapping of the region focused predominantly on white genomes. Further, there’s a paucity of 22q11.2DS diagnosed within Black populations. Thus, there is a requirement to map non-white genomes and determine the landscape of variants in 22q11.2 in genomes with different backgrounds. However, despite the existence of several population groups as publiclyavailable samples, previous studies in Chileans, which are admixed individuals, revealed genetic differences from other population groups [14]. The present study limited the groups of comparison to Black and white genomes.

Given the complexity of the segmental duplications (SDs) residing in the 22q11.2 region, coupled with the high variation in SD copy number, the most accurate approach to resolving the region has been via optical mapping-based methods [9, 10]. Here we optically mapped unaffected and affected Black genomes and compared them to the more widely mapped white genomes to find possible structural variants that are more permissive to the recombination event (NAHR) leading to a deletion in 22q11.2. In unaffected individuals, vastly different haplotype configurations between the 2 groups were discovered with significantly more unique configurations private to Blacks. Further, analysis of the shared configurations revealed a vastly different distribution of haplotypes as well. Additionally, orientation of SD22 modules and spatial locations within a haplotype were also different. Finally, only Black genomes had LCR22D haplotypes with SD22-4 inversions. Altogether, this demonstrated haplotype configurations private to the 2 groups, highlighting the strengths of optical mapping different population groups beyond the often-analyzed white genomes.

The effect of different haplotype configurations ultimately led to different NAHR scenarios. We found most NAHR events to be private to Black PoDOs and white PoDOs. Previous studies found inter-LCR distances and lengths of homology as features which impact NAHR in various diseases stemming from microdeletions [12]. We found that normal Black genomes had more haplotypes which led to shorter lengths of homology and longer inter-LCR22 distances, which are postulated to help explain the decreased incidence of NAHR in Black patients. However, Black PoDOs had NAHR events involving inverted SD22 modules in LCR22A and notably in LCR22D. The implications of this are that the normal genomes’ haplotype frequencies, in either group, directly affect which haplotypes will undergo NAHR. Perhaps these features are not relevant to 22q11.2DS on a per-population basis or they must be re-configured to meet the standards of haplotypes present in each group. However, our small sample size means additional Black PoDOs and probands must be mapped before making definitive conclusions on the features affecting NAHR.

While no whole configuration not present in normal genomes could be pinpointed to be directly responsible for NAHR, these results demonstrate the requirement to map different groups and analyze NAHR scenarios on a per-group basis. When more genomes are mapped, the production of population-specific maps and NAHR permissive events may emerge. More genomes of each group and other populations are currently being mapped toward this goal. Finally, since optical mapping can resolve complex haplotypes in genomic regions intractable to sequencing, these other regions will eventually be mapped and analyzed with population-specific maps as a database.

Overall, this study demonstrated the power of optical mapping of a complex region and the importance of using samples from different backgrounds. In the future, more samples need to be optically mapped across different populations and supplemented with new long read sequence technologies to provide the potential for complete end-to-end haplotypes. In addition, as long-read sequencing technologies become more cost-effective, perhaps the sequence motifs may be studied to provide a lexicon of potential NAHR permissive configurations.

## 5. Acknowledgements

These studies were supported in part by Funds from the NIH grant GM125757 (BSE) and the Charles E.H.Upham endowed chair (BSE).

